# Automated prediction and annotation of small proteins in microbial genomes

**DOI:** 10.1101/2020.07.27.224071

**Authors:** Matthew G. Durrant, Ami S. Bhatt

**Affiliations:** Department of Medicine (Hematology, Blood and Marrow Transplantation) and Department of Genetics, Stanford University, Stanford, California, USA

## Abstract

Recent work performed by Sberro et al. (2019) revealed a vast unexplored space of small proteins existing within the human microbiome. At present, these small open reading frames (smORFs) are unannotated in existing reference genomes and standard genome annotation tools are not able to accurately predict them. In this study, we introduce an annotation tool named *SmORFinder* that predicts small proteins based on those identified by Sberro et al. This tool combines profile Hidden Markov models (pHMMs) of each small protein family and deep learning models that may better generalize to smORF families not seen in the training set. We find that combining predictions of both pHMM and deep learning models leads to more precise smORF predictions and that these predicted smORFs are enriched for Ribo-Seq or MetaRibo-Seq translation signals. Feature importance analysis reveals that the deep learning models learned to identify Shine-Dalgarno sequences, deprioritize the wobble position in each codon, and group codons in a way that strongly corresponds to the codon synonyms found in the codon table. We perform a core genome analysis of 26 bacterial species and identify many core smORFs of unknown function. We pre-compute small protein annotations for thousands of RefSeq isolate genomes and HMP metagenomes, and we make these data available through a web portal along with other useful tools for small protein annotation and analysis. The systematic identification and annotation of those important small proteins will help researchers to expand our understanding of this exciting field of biology.

## Introduction

Small open reading frames (smORFs; ≤50 amino acids in length) and the small proteins they encode play important roles in microbes, including housekeeping, phage defense, and cell signaling functions. These ‘microproteins’ have been identified across multiple domains of life, and given their potential role in mediating cell-cell communication, have been the topic of growing interest in various fields of biology and translational medicine (Hanada et al., 2013; Leslie, 2019; Makarewich et al., 2018; Storz et al., 2014). Despite their importance, the small open reading frames (smORFs) that encode these proteins are difficult to identify, and as a result they are often overlooked and lack functional annotation (Duval & Cossart, 2017; Storz et al., 2014; Su et al., 2013). Techniques such as ribosome profiling (Ribo-Seq) and proteomic approaches have had some success and provide strong evidence of transcription and translation of candidate smORFs (Aspden et al., 2014; Lohmann et al., 2020; Miravet-Verde et al., 2019; Weaver et al., 2019). However, these approaches are limited by a considerable experimental bottleneck, usually requiring the isolation and culture of an organism of interest, and they can only detect what is being actively translated. Thus, Ribo-Seq annotation of smORFs and small protein detection by proteomics approaches are limited in their application.

Microbial smORFs can be difficult to detect using computational annotation of sequenced genomes due to their small size, which makes it difficult to distinguish true smORFs from spurious ones (Hyatt et al., 2010). In the past, many small proteins were discovered largely by serendipity, either overlapping noncoding RNAs or in the intergenic spaces between large ORFs (Jørgensen et al., 2013; Pinel-Marie et al., 2014). A more systematic way to identify and annotate smORFs within microbial genomes is thus very much needed.

Recently, a bioinformatic analysis used evolutionary conservation signals to enhance smORF prediction and thereby accurately identified thousands of small protein families in human-associated metagenomes at a large scale (Sberro et al., 2019). Unfortunately, many of these small open reading frames (smORFs) remain unannotated in existing microbial reference genomes and standard genome annotation tools do not accurately predict them. This limits researchers who focus on specific microorganisms from designing experiments geared toward understanding the role of specific smORFs in the biology of these organisms. For example, annotating the smORFs may offer researchers who have performed dense transposon mutagenesis screens in bacteria an opportunity to connect interesting phenotypes to previously unannotated small genes. While the small protein families identified by Sberro et al. provide a larger set of candidate proteins, no method exists to automatically annotate smORFs in existing genomic sequences from bacterial isolates. Some recent studies have shown that logistic regression and support vector machines (SVMs) hold promise as methods to identify small proteins (Lei Li & Chao, 2020; Zhu & Gribskov, 2019). In one case, the source code is not yet available and it has not yet been published in a peer-reviewed journal, making it difficult to evaluate and understand the underlying approach. In the other case, the model’s precision on bacterial micropeptides from outside of their training set was not assessed. Being able to annotate and identify these small proteins in microbial genomes is an important step toward understanding these proteins and their diverse functions. To that end, a computational tool that can streamline their annotation and can easily be applied to any sequenced genome or metagenomes would be an important step toward understanding the biological functions of these microproteins.

Using the >4,500 small protein families identified by Sberro et al. (2019), we sought to build an annotation tool that combines profile Hidden Markov models (pHMMs) and deep learning models to annotate small proteins in genome and metagenome assemblies. Deep learning has rapidly increased in popularity in the field of genomics, and has been applied to the task of ORF prediction generally (Al-Ajlan & El Allali, 2019). Deep learning models obviate the need for “feature engineering”, the practice of summarizing raw features into metrics and statistics that are believed to be more predictive, and can learn important features automatically by analyzing raw sequence data itself (Zou et al., 2019). However, deep learning often does require a careful model architecture and hyperparameter selection process to achieve optimal performance (L. Li et al., 2017), and this process can be computationally expensive.

Here, we develop *SmORFinder*, an annotation tool that combines profile HMM and a neural network classifier to identify small ORFs in microbial genomes. First, we train deep learning models that analyze the predictions of the Prodigal ORF annotation tool (Hyatt et al., 2010) to determine if the predictions are true smORFs, using the Sberro et al. (2019) data as a training set. We demonstrate that the deep neural networks are more sensitive and have a higher F1 score than profile HMMs when it comes to the classification of protein families that did not exist in the training set. Then we apply this annotation tool to Ribo-Seq and MetaRibo-Seq (Fremin et al., 2020) datasets, demonstrating that its predictions are enriched for actively translated genes, and that combining predictions from different models improves precision. Next, we find evidence that the deep neural networks we trained have learned to identify Shine-Dalgarno sequences, to deprioritize the wobble position in each codon, and to group codons in a way that strongly corresponds with the codon table. Finally, we re-annotate all genomes in the RefSeq database, making the standalone tool and annotations freely available to the research community.

## Results

### Deep learning models detect unobserved smORF families with greater recall and F1 score than profile HMM models

Profile Hidden Markov Models (pHMMs) are widely used in bioinformatics to annotate proteins that are believed to belong to a certain family, or to contain specific domains (Eddy, 1998). Annotation tools such as Prokka (Seemann, 2014) use pHMMs built to recognize specific protein domains which can then be used to annotate predicted microbial ORFs. We sought to compare deep learning models to pHMMs for predicting smORFs. To maximize the potential for comprehensive annotation of smORFs, including those that are highly divergent from those included in the training set, we optimized for deep learning models that performed well on smORF families that were intentionally excluded from the training set (Fig. S1). This gives us estimates for how well a deep learning model is able to generalize to smORF families not seen in the training set (unobserved smORFs) relative to pHMMs.

We used a model architecture that took three different nucleotide sequences as inputs - the smORF itself, 100 bp immediately upstream of the smORF, and 100 bp immediately downstream of the smORF. Using our training set of predicted true positive smORFs (positives) and predicted true negative smORFs (negatives), we chose to tune hyperparameters using the hyperband algorithm (L. Li et al., 2017) to identify two deep learning models. The first model was tuned to have the lowest validation loss on a holdout set of observed smORF families, and the second model was tuned to have the highest F1 score on a holdout set of unobserved smORF families. The three input sequences to each model were handled as three separate branches into one-dimensional convolutional neural networks (CNN), with hyperparameters such as the number of layers in each branch and the filter sizes being tuned independently for each branch. Additionally, we tuned the same model but with bidirectional long short-term memory (LSTM) layers (Tavakoli, 2019) added to the end of each individual branch, with a variable number of neurons in the layer. The three branches are then flattened and concatenated, processed by a single “Dense” layer, and then a final output layer with a sigmoid activation function. We refer to these deep learning model architectures generally as DeepSmORFNets (DSN). We settled on one model that was found to minimize the observed smORF family loss (DSN1), and one model that minimized the unobserved smORF family F1 score (DSN2) (Fig. S2). The models differ in interesting ways: of note, DSN2 uses a bidirectional LSTM layer while DSN1 does not, and DSN1 has more convolutional layers and fewer total parameters (Fig. 1b).

**Figure 1.**
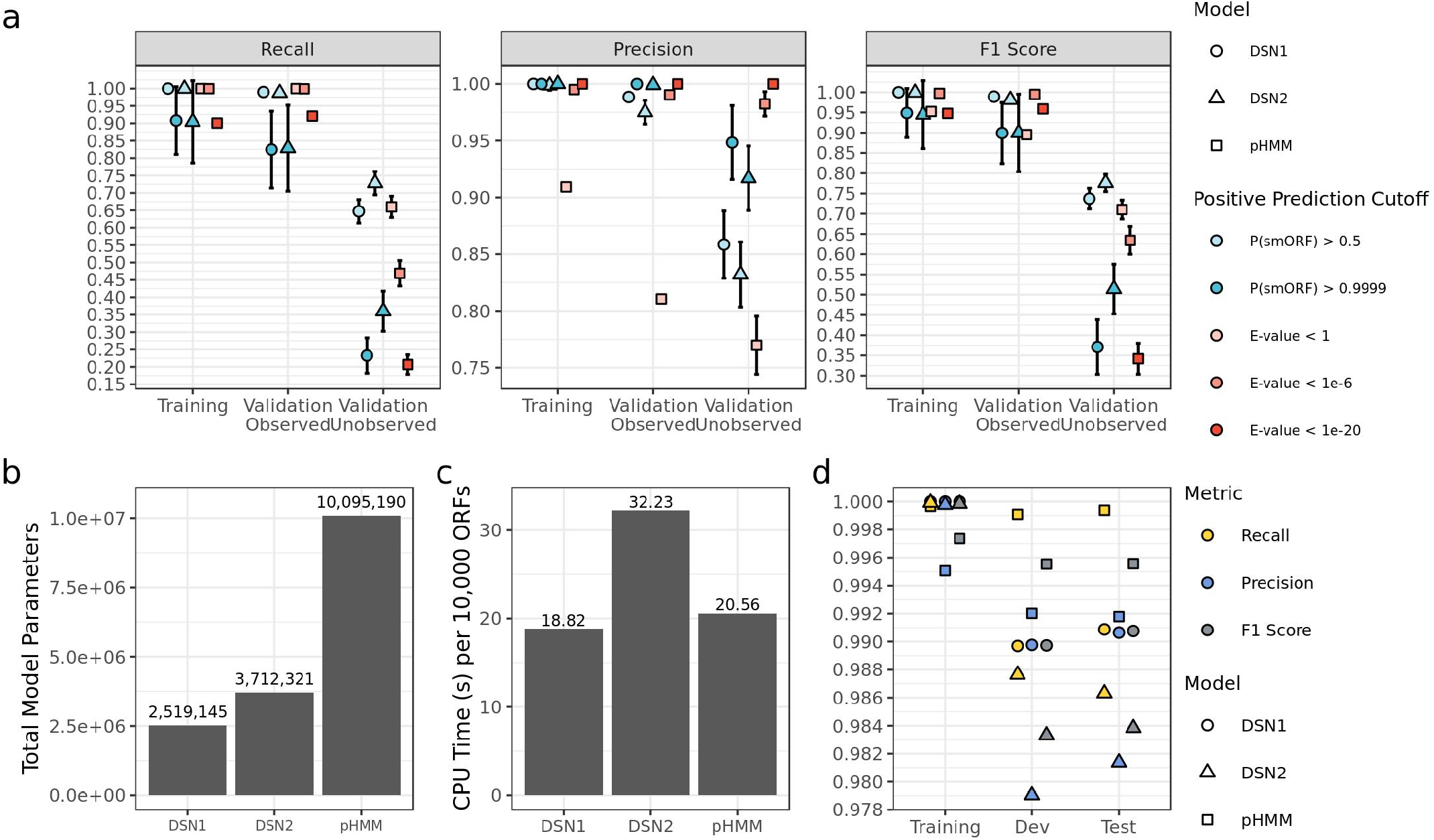
Deep learning models detect unobserved smORF families with greater recall and F1 score than profile HMM models. **a**, Each point is the average value after running the training procedure 64 times with randomly selected families excluded from the training set and assigned to the “Validation - Unobserved Families” set. Showing F1 Score (the weighted average of precision and recall), Precision, and Recall for the three sets with different significant cutoffs. Error bars are the standard error of the 64 randomized samples. **b**, Comparison of the total parameters in the finalized DSN1, DSN2, and pHMM models. **c**, Comparison of the CPU time to process 10,000 ORFs across DS1, DSN2, and pHMM models. **d**, The F1 Score, Recall, and Precision of the final DSN1, DSN2, and pHMM models. A positive prediction cutoff of P(ORF) > 0.5 was used for DSN, and a cutoff of E-value < 1e-6 was used for pHMM.

We then compared the performance of these models against pHMMs. We combined the training and validation (a set not used for training the model, but was used to pick a final model architecture) sets into a single set, and then randomly split into training and validation sets, taking care to ensure that approximately 50% of each validation set consisted of unobserved smORF families (smORF families that did not exist in the corresponding training set). Each random split resulted in a unique training set, a validation set of observed smORF families (Validation - Observed), and a validation set of unobserved smORF families (Validation - Unobserved). We repeated this process 64 times to create 64 randomly split datasets, and chose the model with the lowest loss in the observed smORF validation set. Likewise, we trained pHMMs on the 64 randomized training sets, and compared their performance to the deep learning models on the validation set (Fig. 1).

We find that DSN1, DSN2, and the pHMMs all perform well on the “Validation - Observed” set (Fig. 1a), with Recall, Precision, and F1 Scores that exceed 0.975 for both DSN1 and DSN2 at P(smORF) > 0.5, and 0.99 for pHMMs at E-value < 1e-6. Comparing the performance on “Validation - Observed” set indicates little advantage to using a deep learning model over pHMMs. However, performance on the “Validation - Unobserved” set shows a clear difference between deep learning models and pHMMs. At a cutoff of P(smORF) > 0.5, DSN2 has better average recall than pHMMs at a lenient cutoff of E-value < 1 (Paired t-test; P < 1 x 10^-16^) and better F1 score (Paired t-test; P < 1 x 10^-16^). At the same cutoff, DSN1 has slightly worse recall than pHMMs at a lenient cutoff of E-value < 1 (Paired t-test; P = 0.00684), but a slightly better F1 Score (Paired t-test; P = 1.9 x 10^-10^). These results suggest that the deep learning models are better at generalizing to the unobserved smORF families overall, while the precision of pHMMs continues to be superior at significance cutoffs below 1 x 10^-6^. This suggests the two models may complement each other when used together to identify smORFs.

Despite the increased precision of pHMMs, there are several advantages to the deep learning models., Both deep learning models have fewer learnable parameters than the pHMMs (Fig. 1b), with the pHMMs having 4x more parameters than DSN1. As implemented in python’s keras package and the command line tool hmmsearch, DSN1 runs about as fast as the pHMMs, while DSN2 is slower (Fig. 1c) (Chollet & Others, 2015; *HMMER*, n.d.). Notably, the deep learning models require no sequence clustering or alignment, just the raw smORF and flanking nucleotide sequences. To construct pHMMs for each smORF family, they must be clustered and aligned, with a different pHMM being built for each family. This alignment-free approach to building the model can be considered an advantage of the deep learning models. However, the pHMMs are generative models and require no negative examples for training, while the deep learning models do require these negative examples.

We trained the models one final time on the final training set, which includes at least one representative of each smORF family. We then evaluated all three models on a test set that was held out entirely from the hyperparameter tuning process. We find that all three models perform well on the validation and test sets, with the pHMMs performing the best overall, but all three models having recall, precision and F1 scores that exceed 0.98 on the test set (Fig. 1d).

### Predicted smORFs are enriched for Ribo-Seq signal

We next sought to determine how the models performed on unobserved experimental data. We decided to gauge the quality of the smORF predictions using Ribo-Seq signal as a proxy. Ribo-Seq, a method for ribosome profiling, can identify mRNA sequences that are directly bound by a ribosome, indicating active translation (Ingolia et al., 2009). We reasoned that a more accurate set of smORF predictions would be more likely to be translated, and thus more likely to be enriched for a Ribo-Seq signal. We generated and sequenced Ribo-Seq libraries for one *Bacteroides thetaiotaomicron* isolate (previously published (Sberro et al., 2019)), and four metagenomic human microbiome samples using MetaRibo-Seq, a technique for metagenomic ribosome profiling (Fremin et al., 2020).

We packaged all three models (pHMMs, DSN1, and DSN2) together into a single command line tool that we refer to as *SmORFinder*. We kept all predicted smORFs that met a pHMM cutoff of E-value < 1.0, a DSN1 cutoff of P(smORF) > 0.5, or a DSN2 cutoff of P(smORF) > 0.5. We found 15 smORF families were false positives, and they are automatically excluded from consideration by *SmORFinder*. These families corresponded to the N-terminus of Peptide chain release factor RF2 (prfB) in many different species. This gene contains a naturally occurring programmed frameshift that is corrected upon translation (Curran, 1993), and the Prodigal annotation tool fails to account for this, leading to a spurious smORF annotation.

In *B. thetaiotaomicron*, we find that 86.1% of non-smORFs (ORFs coding for proteins greater than 50 aa in length) have a Ribo-Seq signal (Reads Per Kilobase Million (RPKM) ≥ 0.5; Fig. 2a). In this bacterium, the smORFs tend to have a lower proportion of Ribo-Seq signal, with 86.1% of non-smORFs meeting the RPKM cutoff compared to 23.6% of smORFs. We find that genes predicted by at least one model to be true smORFs are more likely to be enriched for Ribo-Seq signal than rejected smORF predictions. The set of predicted smORFs that met stringent significance cutoffs for DSN1, DSN2 and pHMMs was found to be significantly enriched for Ribo-Seq signal over the smORFs that were rejected by all three models (Fisher’s exact test; P=0.0250). This set was identical to the set identified by DSN1 alone with a high significance cutoff of P(smORF) > 0.9999. The small number of predicted smORFs in this bacterium reduces the power to detect a strong Ribo-Seq enrichment.

**Figure 2.**
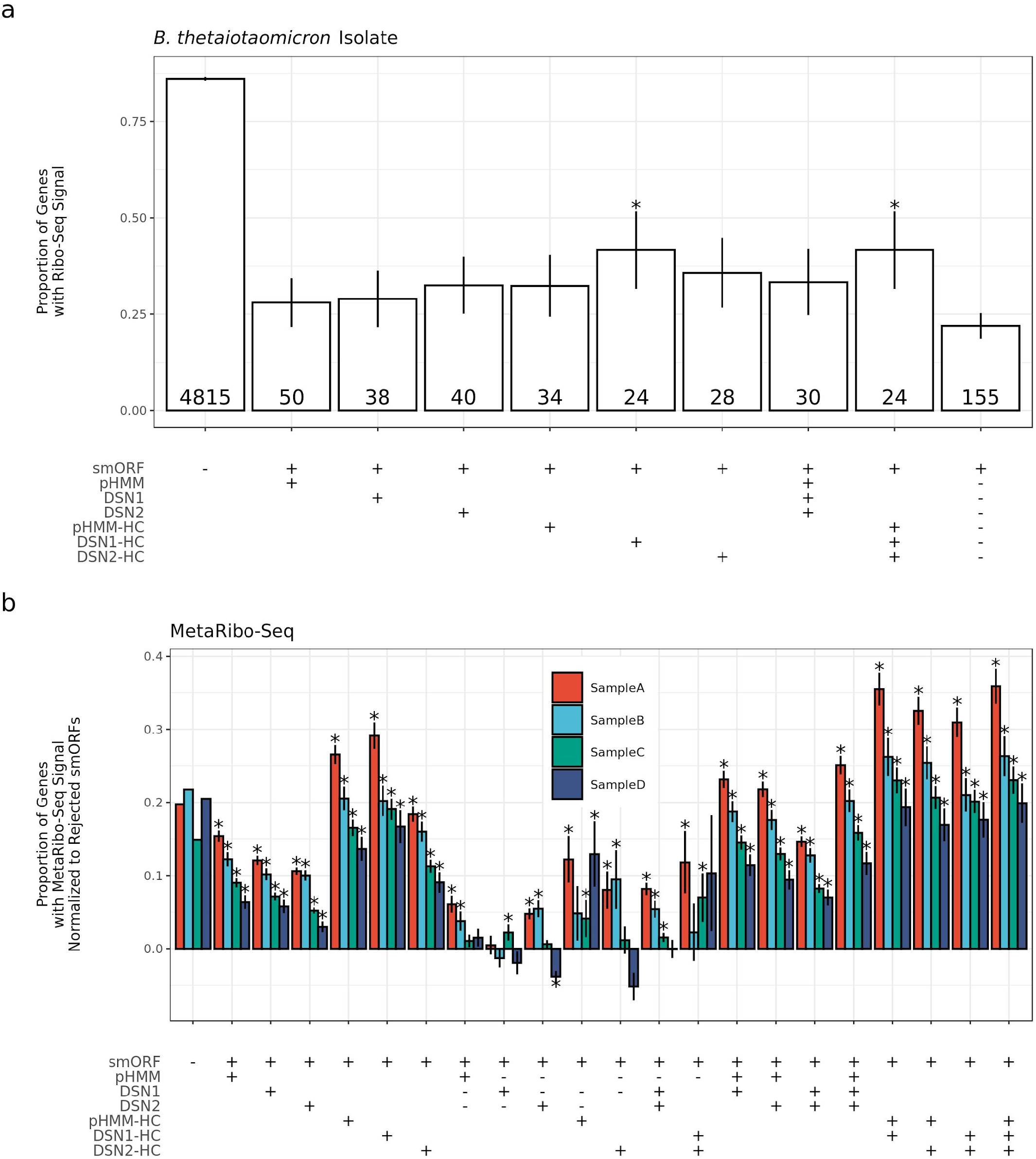
Predicted smORFs are enriched for Ribo-Seq signal. **a**, The proportion of genes with Ribo-seq signal (RPKM ≥ 0.5) in different gene sets in a *Bacteroides thetaiotaomicron* isolate. Table along the x-axis denotes the genes included in each set. The label “smORF” indicates if set includes smORFs (+) or only non-smORFs (-), “pHMM” indicates that the set includes pHMM-predicted smORFs at E-value < 1.0 (+), “DSN1” indicates the set includes DSN1-predicted smORFs at P(ORF) > 0.5 (+), “DSN2” indicates the set includes DSN2-predicted smORFs at P(ORF) > 0.5 (+), “pHMM-HC” indicates the set includes pHMM-predicted smORFs at E-value < 1e-6 (+), “DSN1-HC” indicates the set includes DSN1-predicted smorfs at P(ORF) > 0.9999, and “DSN2-HC” indicates the set includes DSN2-predicted smorfs at P(ORF) > 0.9999. If multiple “+” symbols are found in a column, then that means all genes in the set meet each cutoff. The symbol indicates that all genes meeting the specified cutoff were excluded from the set. The final column indicates all smORFs that were not predicted by any model to be a true smORF at any significance cutoff. Error bars indicate the standard error of each proportion. These smORFs are referred to as “Rejected smORFs”. The number of total genes in each gene set is given at the bottom of each bar. Asterisks indicate that the proportion is significantly higher (P < 0.05) in the specified set than in the “Rejected smORFs” set. **b**, The proportion of genes with MetaRibo-seq signal (RPKM ≥ 0.5), normalized to rejected smORF MetaRibo-Seq signal, in different gene sets in four different MetaRibo-seq samples. Normalization was performed by subtracting the proportion of rejected smORFs with a MetaRibo-Seq signal from the proportion of genes in each set with MetaRibo-Seq signal. The x-axis table is the same as the one shown in (**a**) with additional gene sets added. For example, one additional column is the 8th column from the left, designated by “smORF = +”, “pHMM = +”, “DSN1 = -”, and “DSN2 = -” indicates the set of smORFs with predicted by the pHMM model to be a true smORF, but predicted by both DSN1 and DSN2 to be a false smORF. The gene set with the highest average MetaRibo-Seq symbol across all four samples is the final column on the right, which includes all smORFs that meet stringent significance cutoffs for all three models. Asterisks indicate that the proportion is significantly higher (P < 0.05) in the specified set than in the “Rejected smORFs” set.

We repeated this analysis using published MetaRibo-Seq data generated from stool samples of four human subjects (Fig. 2b) (Fremin et al., 2020). Given the high number of smORFs in each sample, there is increased power to detect a MetaRibo-Seq enrichment compared to the previous example when we only queried a single microbial strain (*B. thetaiotaomicron*). We find in general that predicted smORFs are much more enriched for MetaRibo-Seq signal than “Rejected smORFs” (smORFs that did not meet minimum significance cutoffs for any of the three models). In these metagenomic samples, the MetaRibo-Seq enrichment of predicted smORFs even exceeds the non-smORF enrichment for high-confidence sets that meet high significance thresholds for one or more of the models. As we require higher significance thresholds for the three models (pHMM, DSN1, DSN2), the MetaRibo-Seq enrichment increases across the four samples. The set with the greatest MetaRibo-Seq enrichment over the “Rejected smORFs” is the one that requires high-confidence significance scores across all three models. This suggests that there is value in combining the predictions of the three sets to get a highly precise set of smORF predictions.

### Feature importance analysis reveals inner workings of deep learning models

Deep learning models have been criticized in the past due to their lack of interpretability, often described as a “black box”. Recent advances in deep learning interpretation have overcome this challenge, enabling us to gain insight into the features of the input that play a role in the final model prediction, a technique called feature importance analysis (Shrikumar et al., 2017). We applied feature importance analysis to our deep learning models using the Deep Learning Important FeaTures (DeepLIFT) method as implemented in the SHapley Additive exPlanations (SHAP) python package (Lundberg & Lee, 2017). Briefly, this method calculates the importance of individual input features relative to a set of randomized references by backpropagating the contributions of all neurons to every feature of the input. In the case of our smORF nucleotide sequences, this results in importance scores (also called contribution scores) assigned to each nucleotide in the sequence. For example, if a deep learning model were built to identify ChIP-seq binding sites for a given transcription factor, such as CTCF, feature importance analysis using the DeepLIFT method would identify the CTCF binding motif in individual examples, producing experimentally actionable information.

We apply this technique to both DSN1 and DSN2 to see if we can gain insight into how the two models identify true smORFs. First, we analyze the average DeepLIFT importance scores of all upstream and downstream smORF-flanking nucleotide sequences found in the training set (Fig. 3a). In the upstream sequence, we see a distinct peak at −12 bp in both the DSN1 and DSN2 importance scores. This is in the range of where we typically find the Shine-Dalgarno sequence (a well-described and conserved ribosomal binding site), and upon inspection of individual examples we see that both models did in fact identify the AGGAGG Shine-Dalgarno motif as informative discriminating features (Fig. S3). Interestingly, DSN2 places higher importance on the upstream sequence, perhaps indicating why DSN2 has greater generalizability than DSN1. In the downstream sequence, it would appear that DSN1 places greater importance on the more proximal nucleotide sequences, while DSN2 seems to have identified two particularly important positions at +13 and +4 positions downstream from the stop codon of the smORF. At positions +1 through +10, the nucleotides with the highest average importance scores for DSN2 are all adenine, with the exception of position +3 which is a thymidine. At positions +10 through +20, the nucleotides with the highest average importance scores for DSN2 are all adenine. This suggests that DSN2 has determined that an A-rich downstream sequence may be predictive of a true smORF. By contrast, DSN1 places greater importance on cytosines in the downstream sequence, although it assigns much less importance to the downstream sequence overall. The prioritization of A-rich downstream regions may indicate the rho-independent (intrinsic) transcription termination mechanism, which includes a chain of uracils in the mRNA transcript (d’Aubenton Carafa et al., 1990; Peters et al., 2011).

**Figure 3.**
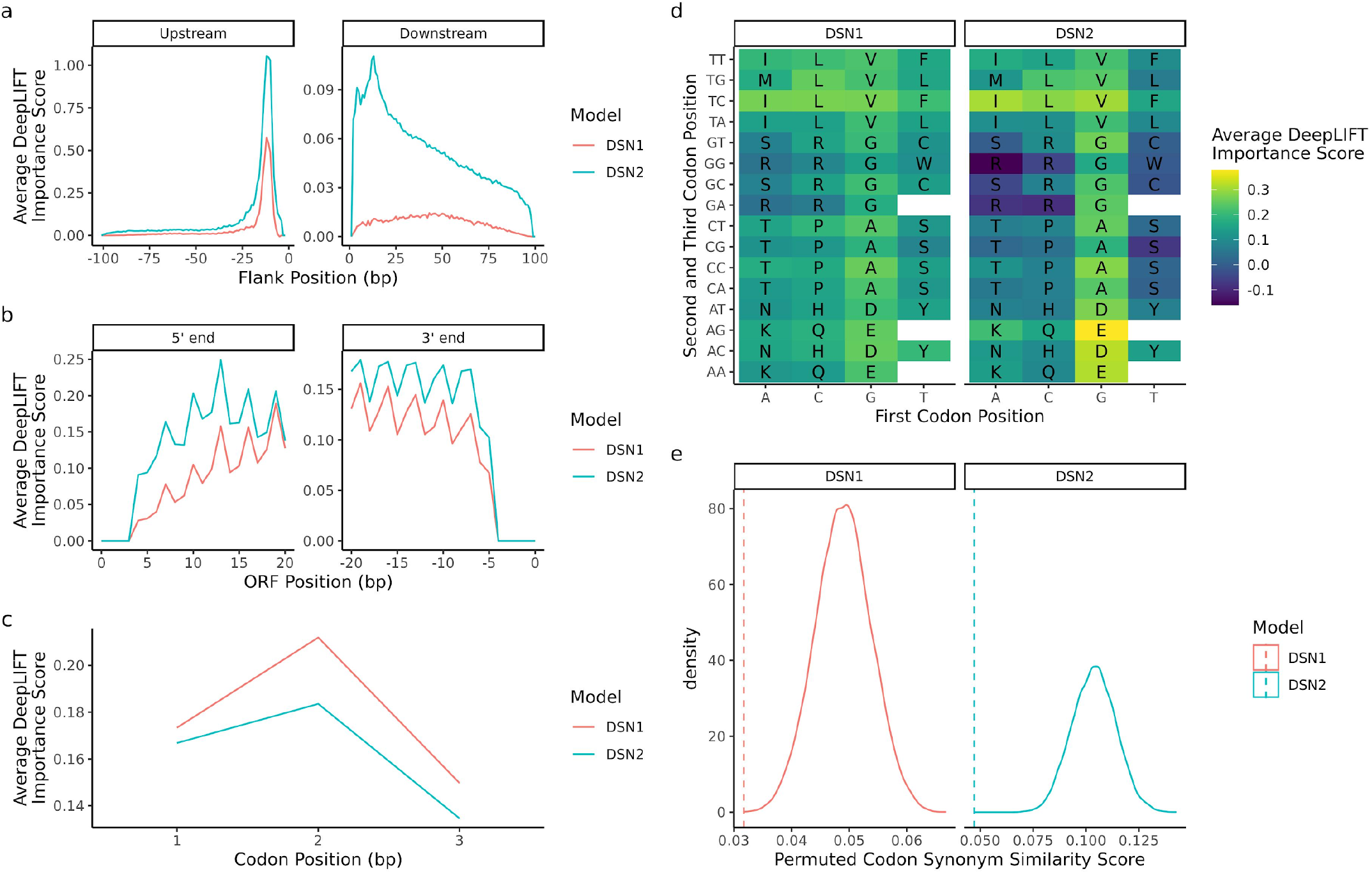
Feature importance analysis reveals inner workings of deep learning models. **a**, Average feature importance scores across the 100 bp upstream and downstream of each true smORF example in the training set. Showing the feature importance scores for DSN1 (red) and DSN2 (blue). **b**, The average feature importance scores across the first 21 and last 21 bp of all true smORF examples in the training set. **c**, The average feature importance scores of the first, second, and third codon position in codons of each true smORF example in the training set. Both models assign higher feature importance to the first two codon positions than the third (wobble) position. **d**, The average feature importance scores of each codon in the codon table, excluding stop codons. The nucleotide of the first codon position is on the x-axis, while the second and third positions are shown on the y-axis. **e**, The true codon synonym similarity (CSS) score (dotted lines) vs. the distribution of CSS scores (solid line) observed when randomly permuting codon synonym labels across all scores.

Next, we analyze the average importance scores across the first 21 and the last 21 base pairs within each smORF (Fig 3b). The base pairs found in the initiation codon and stop codon have importance equal to zero because those regions were not permuted in the randomly generated reference sequences. There appears to be some difference across both scores for the two models, but what is most striking is the obvious periodicity in the signal. This is not surprising considering the periodic nature of codons found in functional ORFs. When we average the importance scores across all codons, we see that both DSN1 and DSN2 place greatest importance on the second codon position, and the least importance on the third codon position or “wobble” position (Fig. 3c). While it is not clear why greater importance would be placed on the second codon position compared to the first codon position, the fact that the wobble position has less overall importance is intriguing considering its redundant role in the codon. This implies that the model has learned to identify these codons and to give lesser importance to the wobble position when making its predictions.

We next look at average importance scores of each unique codon across all true smORFs (Fig. 3d). We find that codons are highly correlated in their average importance across the two models (Spearman r = 0.906; P < 2.2 x 10^-16^; Fig. S3). When the importance score of each amino acid is averaged across all codon synonyms, glutamate, aspartate, valine, and alanine have the four highest average importance scores for both DSN1 and DSN2. The four amino acids with the lowest average importance scores across the two models are arginine, serine, tryptophan, and cysteine. In the case of DSN2, the average importance scores of these four amino acids are actually negative, indicating that on average these amino acids actually prompt the model toward making negative predictions, implying that in certain contexts true smORFs may typically lack these amino acids.

Finally, we investigated if the deep learning models learned to assign similar importance to codon synonyms. This implies that some representation of the codon table was learned during training. We developed a Codon Synonym Similarity Score (CSS score), which is the average standard deviation of importance values among codon synonyms (Fig. 3e). We first calculated the CSS score for the DSN1 and DSN2 models, and then we permuted the codon synonyms across the importance scores to get a null distribution of CSS scores. We find that for both models, the true CSS score is very low in the range of permuted CSS scores, indicating that codon synonyms share similar importance scores, and that some representation of the codon table was learned by the model.

### Core genome analysis identifies core smORFs of unknown function

Seeing the value in pre-computing smORF annotations for RefSeq genomes for the scientific community, we used *SmORFinder* to analyze 191,138 RefSeq genomes, in addition to the HMP metagenomic samples that were used as part of the initial smORF family identification process. This included genomes across 63 bacterial phyla, with 104,658 genomes belonging to members of the Proteobacteria phylum, and 19,681, 12,338, and 11,511 genomes belonging to the *Escherichia coli, Staphylococcus aureus*, and *Salmonella enterica* species, respectively. These data, along with other useful tools for smORF analysis, are available through our web portal (Fig. S4) that can be accessed through our github repository at https://github.com/bhattlab/SmORFinder.

We carried out a core-genome analysis of 26 of the most common species’ genomes found in RefSeq (Table S1). Genomes were included only if they had an average Mash (Ondov et al., 2016) distance that exceeded 0.95 with all other genomes in the set, ensuring that the genomes were related enough to each other in each collection to be considered the same “species”. We clustered genes at 80% identity using CD-HIT, and identified genes that were found in the core genome (present in greater than 97% of all genomes for the species).

We find 692 putative smORFs to be part of the core genome across the 26 species (Fig. 4a). These are all smORFs annotated by Prodigal with lowered minimum size cutoffs, and prior to filtering according to DSN or pHMM significance cutoffs. The total number of such core smORFs varies widely across species, with 106 identified in *Bacillus cereus*, and 4 identified in *Helicobacter pylori*. However, the total number of core smORFs is difficult to meaningfully compare across species, as they vary in their overall diversity. Across all species, 70.7% of these smORFs contain no recognized Pfam domains, 9.54% contain a ribosomal protein domain, and 19.77% contain some other known domain.

**Figure 4.**
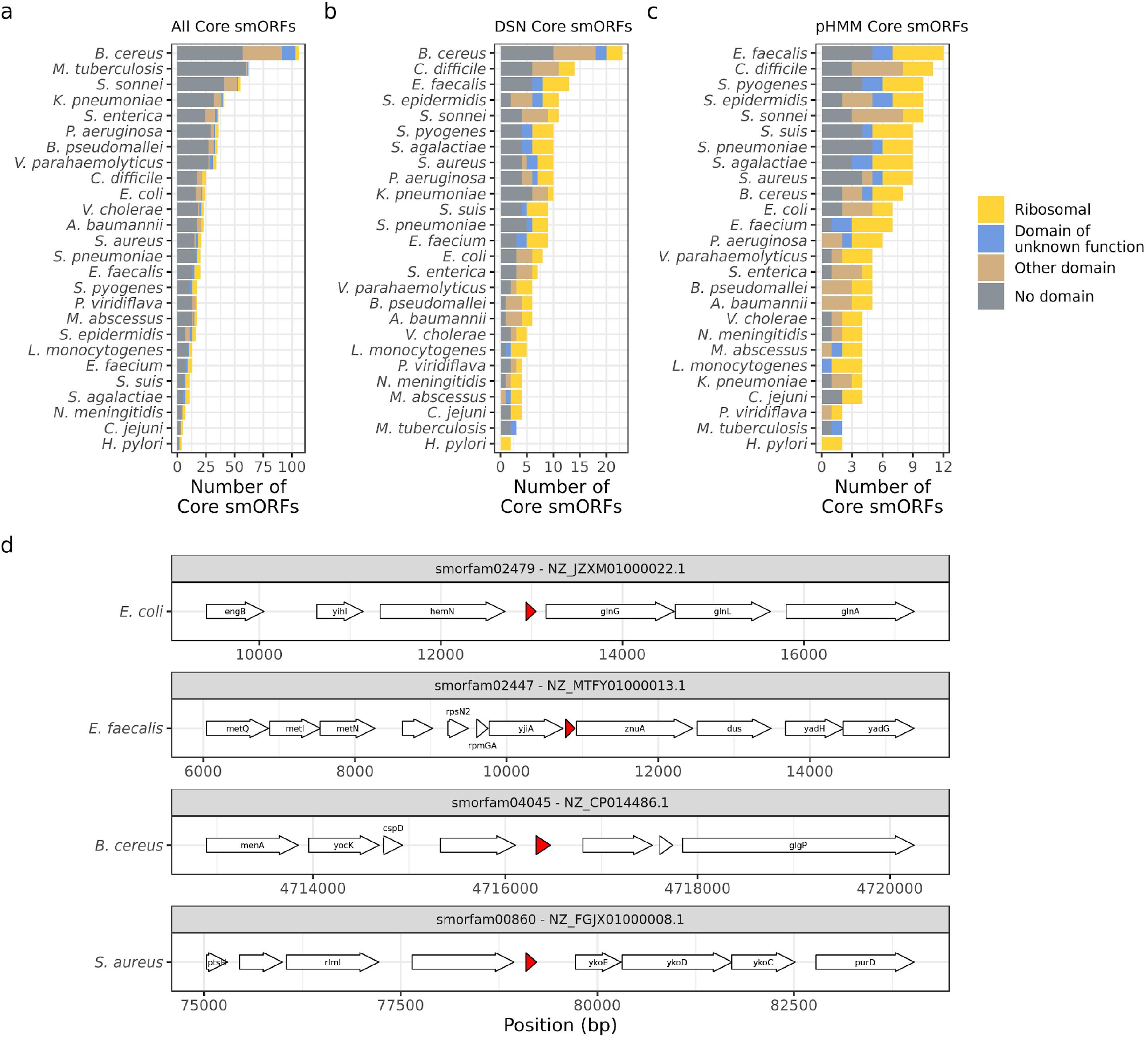
Core genome analysis identifies core smORFs of unknown function. **a**, The total number of core smORFs found in each species’ genomes. These smORFs were not filtered using DSN, all smORFs identified by the Prodigal annotation tool (with a lowered minimum size cutoff of all smORF predictions greater than 15 nucleotides) were included. **b**, The total number of core smORFs identified by the *SmORFinder* (DSN) annotation tool as being true smORFs. This includes all smORFs that meet stringent significance cutoffs for at least one model (pHMM E-value < 1e-6, DSN1 P(smORF) > 0.9999, or DSN2 P(smORF) > 0.9999), or those that meet lenient significance cutoffs for all three models (pHMM E-value < 1, DSN1 P(smORF) > 0.5, and DSN2 P(smORF) > 0.5). **c**, The total number of core smORFs per species that meet the pHMM significance cutoff of E-value < 1e-6. Colors indicate if each core smORF contains a Pfam domain (E-value < 1e-6), and which type. “Ribosomal” (yellow) implies a ribosomal protein domain, “Domain of unknown function” (blue) implies it has a recognized domain of unknown function, “Other domain” (brown) indicates some other Pfam domain, and “No domain” (grey) indicates that it does not contain any known Pfam domain. **d**, Four example core smORFs with no recognized Pfam domain that exist in the core genome of two or more species. Arrows indicate ORFs identified by *SmORFinder* or by the Prokka annotation tool. The red regions indicate the position of each core smORF. The text indicates gene names as assigned by Prokka. The absence of any gene name indicates that Prokka identified the genes as “hypothetical” proteins. The species to which each genome belongs is noted to the left of the gene diagram, the smORF family (smorfam) ID and NCBI Reference Sequence ID are given in the strip above each region.

When we use our DSN predictions as calculated by the *SmORFinder* tool to filter this list of core smORFs, we dramatically reduce the total number from 692 to 213 (Fig. 4b). This enriches for smORFs that contain a predicted Pfam domain, with 31.0% being ribosomal proteins, 30.49% containing some other Pfam domain (including domains of membrane bound YbgT-like proteins, entericidins, and Multidrug efflux pump-associated protein AcrZ among others), and 38.5% containing no Pfam domain. This list can be further reduced by relying only on pHMM models of known smORF families with a stringent significance cutoff (E-value < 1e-6), resulting in 167 such core smORFs (Fig 4c). Using only this significance cutoff as a filter, the total number of core smORFs drops dramatically for some species, such as *B. cereus* whose total number of core smORFs drops from 23 to 8.

We find four smORF families with no recognized Pfam domain that appear in more than two different species’ core genomes (Fig. 4d). The smORF family smorfam02479 is homologous to YshB, a predicted transmembrane protein recently shown to play a role in intracellular replication in *Salmonella* virulence (Bomjan et al., 2019). We find that members of this smORF family exist in the core genomes of other Enterobacteriaceae such as *E. coli, Klebsiella pneumoniae, S. enterica*, and *Shigella sonnei*. The smORF family smorfam02447 shown in Fig. 4d is a 40 aa protein found between P-loop guanosine triphosphatase YjiA and zinc uptake system protein ZnuA in the *Enterococcus faecalis* genome. Members of this smORF family were found in the core genomes of *S. aureus, Streptococcus agalactiae, Streptococcus pyogenes*, and *E. faecalis*, and its function has not been characterized. The smORF family smorfam04045 protein shown in Fig. 4d is a 49 aa protein found between largely uncharacterized protein and a predicted lipase in the *B. cereus* genome. Members of this smORF family were found in the core genomes of *B. cereus, Streptococcus suis, and S. pyogenes*, and it is 91.8% identical to a *Bacillus manliponensis* gene described as an “Alcohol dehydrogenase” in UniProt, although most other homologs are considered to be uncharacterized. The smORF family smorfam00860 shown in Fig. 4d is a 44 aa protein found between an uncharacterized protein and putative HMP/thiamine permease protein YkoE in the *S. aureus* genome. Members of this smORF family were found in core genomes of *S. aureus* and *Staphylococcus epidermidis*, and its function has not been characterized.

## Discussion

Recent advances in bioinformatic annotation approaches and de novo annotation of genes using Ribo-Seq have enabled the discovery of thousands of novel smORFs. The microproteins that they encode have emerged as macromolecules of interest in organisms ranging from microbes to plants to mammals. Unfortunately, to date, no method exists for the accurate annotation of microbial genomes for these smORFs, and most existing microbial genomes are lacking comprehensive annotation for ORFs<150 nucleotides. To overcome this challenge, we present and evaluate the performance of a smORF annotation pipeline. We demonstrate that deep learning models can distinguish between true smORFs and spurious smORFs about as well as pHMMs trained on observed smORF families, and may perform better than pHMMs on unobserved smORF families, suggesting that the model has utility as a de novo smORF identification tool. We find that combining the predictions of pHMMs and deep learning models results in a set of smORF predictions that are enriched for Ribo-Seq translation signals. We acknowledge that in isolation, the deep learning models themselves do not perform significantly better than pHMMs overall. This suggests that pHMM models may already perform at close to the irreducible error of this particular task for observed protein families. Indeed, we can see that for observed protein families, pHMMs already exceed both a recall and precision of 0.99.

We find that both the deep learning models and the pHMMs dramatically increase the Ribo-Seq and MetaRibo-seq enrichment signal of the annotated smORF set. This suggests that selecting smORFs based on the predictions of these models greatly enriches for actively translated and thus likely functional smORFs. Including the three different models (DSN1, DSN2, and the pHMMs) in the *SmORFinder* annotation tool opens up new options for filtering a set of candidate smORFs. For example, rather than relying on a strict significance cutoff for one or multiple models, we find that using lenient significance cutoffs that must be met by all three models is a good strategy for narrowing down a list of candidate smORFs.

New advances in feature importance analysis allows us to peer into the “black box” of deep learning. This is a fascinating look at how these powerful predictive algorithms learn to identify true smORF families, and we can see that they automatically learn features that scientists characterized long ago by experimental means (Shine-Dalgarno sequences, codon periodicity, codon synonyms, etc.). It also acts as an interesting opportunity to find new, generalizable features that may have previously gone unnoticed. For example, DSN2 appears to assign greater importance to 3’ downstream sequences that are A-rich. This could indicate that the model has learned to recognize rho-independent (intrinsic) transcription termination sequences, which are known to contain a chain of uracils in the mRNA transcript (d’Aubenton Carafa et al., 1990; Peters et al., 2011). Intrinsic terminator sequences are not taken into consideration by ORF annotation algorithms such as Prodigal (Hyatt et al., 2010).

Our core genome analysis of 26 different bacterial species identified many smORFs that appear to be highly conserved and possibly essential, which included smORFs that were identified using permissive Prodigal annotation and clustering prior any *SmORFinder* models were applied. Using *SmORFinder* predictions to filter these core smORFs showed a significant reduction in the total number of smORFs for some species. For example, 106 core smORFs were found in *B. cereus* genomes prior to *SmORFinder* filters, and reduced to only 8 core smORFS after applying strict filters. This could indicate that there are many smORFs that were not found in the initial set of core smORFs but are found in the *B. cereus* genome, or that a large number of the core smORFs found in the *B. cereus* genome are false positives. Further experiments and efforts to supplement our set of core smORFs will likely shed light on this question.

While efficient and powerful, the approach that we take in this study has several limitations. First, the *SmORFinder* annotation tool is primarily limited by the Prodigal calling algorithm. The original set of >4,500 families identified by Sberro et al. relied on a downstream analysis of smORFs that were identified by Prodigal with a lowered minimum size threshold. *SmORFinder* is also limited to predictions made by Prodigal, and can be thought of as an additional filter step on top of these Prodigal predictions. Other smORFs that are missed by Prodigal, such as those that significantly overlap with other ORFs, will most likely be missed. Second, we are also limited by the accuracy of the predictions made by Sberro et al. in their original study. In the course of completing this analysis, we identified 15 smORF that are false positives; while this is a relatively small number of overall false positives, it is likely that there are other such false positives in the overall set. Third, the true generalizability of the algorithm we have shown here is also questionable. That is, it does not appear that the model can reliably identify true smORFs that are evolutionarily unrelated to smORFs in the original training set. Our analysis shows that incorporation of the deep learning approaches may improve the identification of distant homologs of smORF families that exist in the training set compared to the pHMMs alone. For many applications, it may be best to only trust predictions that are highly significant, or those that are corroborated by multiple models.

These limitations notwithstanding, with the growing interest in microproteins, *SmORFinder* should be valuable to the research community as it will allow researchers to filter down lists of candidate smORFs to a more accurate list of smORF predictions. We have precomputed the smORFs of thousands of RefSeq isolates genomes and HMP metagenomes and made them available for download through a web portal (Fig. S4). The annotation tool can easily be installed as a python package and is ready for use. This will enable the study of new smORFs, opening up many avenues for biological research. For example, the reannotation of these bacterial genomes could help gain new insights into previously conducted experiments, such as transposon-mutagenesis experiments, affording researchers a wealth of functional data. These data are made freely available for the research community through our github repository and web portal (https://github.com/bhattlab/SmORFinder). It is possible that a suite of tools, including but not limited to SmORFinder, will be developed and applied for the comprehensive, sensitive and specific detection of smORFs across prokaryotes. As such, we anticipate that SmORFinder may be augmented by other models as they are published and thoroughly validated.

## Methods

### Curating positive and negative training examples

A critical first step in any approach to develop a homology/pattern-based annotation algorithm is development of a positive and negative training set. In this case, positive and negative examples of small proteins were needed to train the neural network. Positive examples were derived from the 4,539 small protein families that were originally reported by Sberro et al. (2019). A maximum of 64 examples per protein family were kept, and these 64 were randomly chosen. 100 base pairs of upstream and downstream sequences were used as model inputs, along with the ORF sequence itself. In the event that the upstream and downstream sequences were shorter than 100 base pairs, whatever sequence was available was used. The strategy used to identify negative examples was similar to the one used to identify the positive examples, but with criteria reversed. Prior to applying filters, Sberro et al. identified approximately 400,000 protein family clusters (clustered using CD-HIT with the parameters -n 2 -p 1 -c 0.5 -d 200 -M 50000 -l 5 -s 0.95 –aL 0.95 –g 1). To identify negative examples (small protein families that are most likely spurious ORFs), the following filters were applied. First, smORF clusters were excluded if they were predicted to contain a known protein domain according to an Reversed Position Specific (RPS) BLAST search of the Conserved Domains Database (CDD) database (A predicted domain was considered significant if the e-value was less than or equal to 0.01, and the small protein aligned to at least 80% of the length of the position-specific scoring matrix (PSSM)). Next, families with less than 4 unique members were excluded, as this is too few examples to be properly analyzed by RNAcode. Next, RNAcode was run on each family with the parameter --num-samples 200, and default parameters otherwise. RNAcode can identify conserved coding sequences with samples as small as 4 unique examples, but the average pairwise identity of these examples must be below 90% (Washietl et al., 2011). For families with 4 to 7 members, the RNAcode results were considered only if this average pairwise identity threshold was met for the family. With families that contained >8 members, we required at least one pair to fall below the 90% identity threshold. Any families that were predicted by RNAcode to contain coding sequence (CDS) regions were excluded. Next, protein families were aligned to each other using the DIAMOND search algorithm. If any remaining negative example was a significant (e-value < 1e-3) match for any positive example, it was excluded. If any negative example aligned to another negative example that failed the RNAcode conserved CDS detection, it was also excluded. This resulted in 4,705 high-confidence negative small protein examples. To further supplement this dataset of negative examples, more negative examples were synthesized by shuffling upstream, downstream, and ORF nucleotide sequences. Both positive and negative sequence examples were shuffled using a tetramer shuffling algorithm implemented in the tool by fasta-shuffle-letters in the MEME Suite (Bailey et al., 2009), which was built on the uShuffle algorithm (Jiang et al., 2008). Start and stop codons were preserved, and shuffled ORF sequences were only kept if they could be fully translated using the same translation table as the original sequence. This was to ensure that the deep learning model would learn to discriminate true positives from random sequences. The final ratio of positive to negative examples was 0.417.

### Splitting dataset into training, validation, and test sets

We used a stratified sampling approach to randomly split the full dataset into training, validation, and test sets (Fig. S1). First, we made sure that each dataset had at least one member of each smORF family, which were randomly distributed across the three datasets. After this requirement was met, the remaining examples were randomly allocated to the three different datasets, with approximately 80% being allocated to the training set, 10% to the validation set, and 10% to the test set. The final training set included 367,184 examples (112,427 positive, 254,757 negative), the final validation set included 47,248 examples (13,192 positive, 34,056 negative), and the final test set included 46,932 examples (12,933 positive, 33,999 negative). The training and validation sets were combined and permuted such that certain protein families in the validation set were excluded from the training set (unobserved smORF families). These permuted datasets were used to estimate the performance of the model on the unobserved smORF families.

### Deep learning model architecture and hyperparameter tuning

Hyperparameter tuning was used to identify model architectures that performed best on the validation dataset. The basic model included three inputs, a one-hot encoded vector with dimensions 153×4 to represent the smORF sequence itself, with zeroes padded on the right for smORFs shorter than 153 bp, 100 bp upstream of each smORF encoded as a 100×4 vector, and 100 bp downstream of each smORF encoded as a 100×4 vector. These are then fed into one-dimensional convolutional layers, which are followed by a dropout layer and a pooling layer. All three input branches are flattened and concatenated as a single vector, which is processed by a final dense layer, a dropout regularization, and a final dense layer with a sigmoid activation function that calculates the probability that the input smORF is a true smORF. The many hyperparameters in this model were tuned using the hyperband algorithm (L. Li et al., 2017) as implemented by the keras-tuner python package (O’Malley, 2020). This algorithm randomly sampled the hyperparameter space, including number of convolutional layers per input branch (1, 2, or 3), the number of filters per layer (32, 64, 128, 256, 512, or 1024), the size of each filter (6, 12, 18, or 24), the dropout rate of the convolutional layers (0.1, 0.3, or 0.5), the dropout rate of the final dense layer (0.1, 0.3, or 0.5), the number of neurons in the final dense layer (16, 32, 64, 128, 256, or 512), the learning rate (1e-5, 1e-4, or 1e-3), the padding method (“valid” or “same”), and the pooling method (max pooling or average pooling). Adam optimization with a learning rate of 1e-4 was used to train the model (Kingma & Ba, 2014). After the first convolutional layer, the number of convolution filters is divided by 2, and the filter size is reduced by one-third of the original filter size. For example, if a model had three layers and 1024 filters of length 18, the second layer would have 512 filters of length 12, and the third layer would have 256 filters of length 6. Our aim was to identify models that minimized the loss across the “Validation - Observed” dataset, and maximized the F1 score of the “Validation - Unobserved” dataset. Hyperband was run with a maximum number of epochs of 200 and a downsampling factor of 3 over 4 complete iterations, resulting in 512 different hyperparameter combinations. This was repeated for both the “Validation - Observed” and “Validation - Unobserved” datasets. Finally, the same process was repeated with an additional LSTM layer added at the end of each input branch, which added hyperparameters including the number of LSTM neurons per input branch (16, 32, 64, 128, or 256) and the LSTM dropout rate (0.1, 0.3, or 0.5). DSN1 and DSN2 were chosen as the models with the best loss in calculated over the “Validation - Observed” dataset, and the best F1 score calculated over the “Validation - Unobserved” dataset, respectively. See Fig. S2 for a final description of each model’s hyperparameters.

### Building profile Hidden Markov Models

Using the positive examples (true smORFs) of each protein family found in the training dataset, profile HMM models were constructed. Small protein families were aligned using MUSCLE (Edgar, 2004). We then used the command line tool hmmbuild to construct the pHMM for each family (*HMMER*, n.d.). All pHMMs were combined into a single file, and the e-value of the pHMM with the lowest pHMM is used to assign a given sequence to a smORF to a family.

### Determining how deep learning models and profile HMMs generalize to unobserved smORF families

A permutation approach was used to determine how well the deep learning models and pHMMs generalize to Unobserved smORF families. As depicted in Fig. S1, the training set and validation set were combined, and the two validation sets were created - one that contained smORF families that were observed at least once in the training set (Validation - Observed), and one that contained smORF families that were not observed in the training set (Validation - Unobserved). The smORF families that were excluded from the training set were randomly chosen, and this process was repeated 64 times to create 64 train-validation splits. The deep learning models (DSN1 and DSN2) were both trained on all 64 training sets for up to 2000 epochs to minimize the training loss. Early stopping was used to choose the model that had no improvement in the calculated “Validation - Observed” loss after 100 epochs. This final trained model was then evaluated on the training set and validation sets to estimate the model’s precision, recall, and F1 score. The distribution of these performance metrics across the 64 permuted datasets was used to determine the error of each estimate as shown in Fig. 1a. The pHMM models were also trained independently on the same 64 permuted datasets to get comparable performance estimates.

### Finalizing deep learning model

We trained the final deep learning models and pHMMs on the initial, unmodified training set (Fig. S1). We selected the deep learning models with the lowest loss in the validation set, with a maximum of 2000 training epochs and early stopping after 100 epochs of no improvement in the validation loss. We then evaluated all models on the validation set and the test set, which was held out from the beginning and was not included in the model architecture selection process. These final models are included in the *SmORFinder* annotation tool in its current implementation.

### Validating SmORFinder with Ribo-Seq datasets

Ribo-Seq datasets were used to validate that the *SmORFinder* annotation tool enriches for actively translated smORFs. We used previously published Ribo-seq datasets that are available through the NCBI SRA portal under the projects PRJNA540869 (Ribo-Seq of *B. thetaiotaomicron* isolate) and PRJNA510123 (MetaRibo-Seq of human stool samples from 4 individuals) (Fremin et al., 2020; Sberro et al., 2019). The *B. thetaiotaomicron* reference genome was annotated using Prodigal configured to identify smORFs. Assembled metagenomes of the 4 MetaRibo-Seq samples were also annotated and used as a reference for each respective sample. Ribo-Seq reads were aligned to reference genomes using bowtie2 (Langmead & Salzberg, 2013). Ribo-Seq coverage of each predicted ORF was calculated using bedtools (Quinlan & Hall, 2010). Any ORF that had a calculated RPKM ≥0.5 was considered to have a Ribo-Seq signal. The *SmORFinder* annotation tool was used to identify predicted smORFs. Any smORF that met at least one of the significance cutoffs (pHMM e-value < 1; DSN1 > 0.5; DSN2 > 0.5) was considered a potential smORF. All smORFs that didn’t meet any of these cutoffs were considered “Rejected smORFs”. Different subsets of smORFs were identified based on their statistical significance and agreement across the three different models. These subsets were compared to the “Rejected smORFs” subset in terms of Ribo-Seq signal, and Fisher’s exact test was used to determine if the subset significantly differed.

### Feature importance analysis

A feature importance analysis of both the DSN1 and DSN2 models was performed to interpret, in part, how the deep learning models were learning to identify true smORFs. This was done using the DeepLIFT algorithm (Shrikumar et al., 2017) as implemented in the SHAP python package (Lundberg & Lee, 2017). This technique measures the importance of individual features, nucleotides in this case, in determining the model’s prediction relative to some references. Dinucleotide shuffling of upstream and downstream nucleotide sequences were used as a reference. The start and stop codons of the ORF sequences were preserved in the references, and the intermediate sequence was dinucleotide shuffled until a non-interrupted ORF was generated. Twenty shuffled references were used for each example. Averages across all examples in the training dataset are shown in Fig. 3.

### Codon Synonym Similarity Score

A codon synonym similarity score (CSS score) was calculated to determine how similar the DeepLIFT importance scores were for codons that code for the same amino acid. This was calculated as:

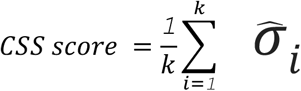

Where 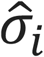 is the standard deviation of the average feature importance scores for codon synonym group *i*. This was calculated for *k* = 18 amino acids, excluding methionine and tryptophan which only have one codon for each. To determine a null distribution of this score, codon synonym labels were randomly permuted across the feature importance scores, and the CSS score was recomputed. This was repeated 10,000 times to calculate a permuted null distribution. The original CSS score was compared to this permuted null distribution to determine its statistical significance.

### Annotating small proteins in RefSeq genomes

All RefSeq bacterial genomes were downloaded on April 29th, 2020. This included all genomes matching the NCBI Entrez search query ‘“Bacteria”[Organism] AND (latest[filter] AND (all[filter] NOT anomalous[filter] AND all[filter] NOT partial[filter]))’. In total, 191,138 genomes were downloaded. These were annotated using the *SmORFinder* annotation tool, and data were compiled into a database that can be accessed through the github repository https://github.com/bhattlab/SmORFinder.

### Core-genome analysis

A core-genome analysis was carried out on 26 bacterial species with a high number of available isolates (Table S1). This included *Acinetobacter baumannii, Bacillus cereus, Burkholderia pseudomallei, Campylobacter jejuni, Clostridioides difficile, Enterococcus faecalis, Enterococcus faecium, Escherichia coli, Helicobacter pylori, Klebsiella pneumoniae, Listeria monocytogenes, Mycobacterium tuberculosis, Mycobacterium abscessus, Neisseria meningitidis, Pseudomonas aeruginosa, Pseudomonas viridiflava, Salmonella enterica, Shigella sonnei, Staphylococcus aureus, Staphylococcus epidermidis, Streptococcus agalactiae, Streptococcus pneumoniae, Streptococcus pyogenes, Streptococcus suis, Vibrio cholerae*, and *Vibrio parahaemolyticus*. Mash distances for all isolates of each species, and only isolates that had an average mash distance < 0.05 (corresponding roughly to 95% average nucleotide identity (ANI)) to all other isolates of the species were kept. All isolate genomes were annotated using Prodigal (Hyatt et al., 2010) to identify all open reading frames, with the minimum gene size filter lowered to 15 nucleotides. All identified protein sequences were then clustered at 80% identity using CD-HIT (Huang et al., 2010), where each cluster represented a unique “gene”. All genes that were found to exist in greater than 97% of all isolates for each species were considered to be part of that species’ “core genome.”

## Acknowledgements

We thank Hila Sberro for assistance with compiling information about the smORFs identified in her original study. We thank Brayon J. Fremin for his help with previously published Ribo-Seq and MetaRibo-Seq datasets and for providing feedback on the manuscript. We thank Soumaya Zlitni, Dylan Maghini, and Chris Severyn for providing feedback on the manuscript.

## Author Contributions

M.G.D. conceived the study and analyses, designed the software, performed formal analyses, visualized the data, wrote the manuscript, and coordinated the project. A.S.B. helped with conceptualization, writing, and editing the manuscript, and funding acquisition.

## Data Availability

The code for *SmORFinder* is available at https://github.com/bhattlab/SmORFinder. The web portal can be accessed through the github repository. The complete training set and validation sets will be made available upon official publication of the manuscript in a peer-reviewed journal. If you want early access to this dataset, please contact the corresponding author.

