## Supplementary Figures for "Automated prediction and annotation of small proteins in microbial genomes"

### Supplementary Materials

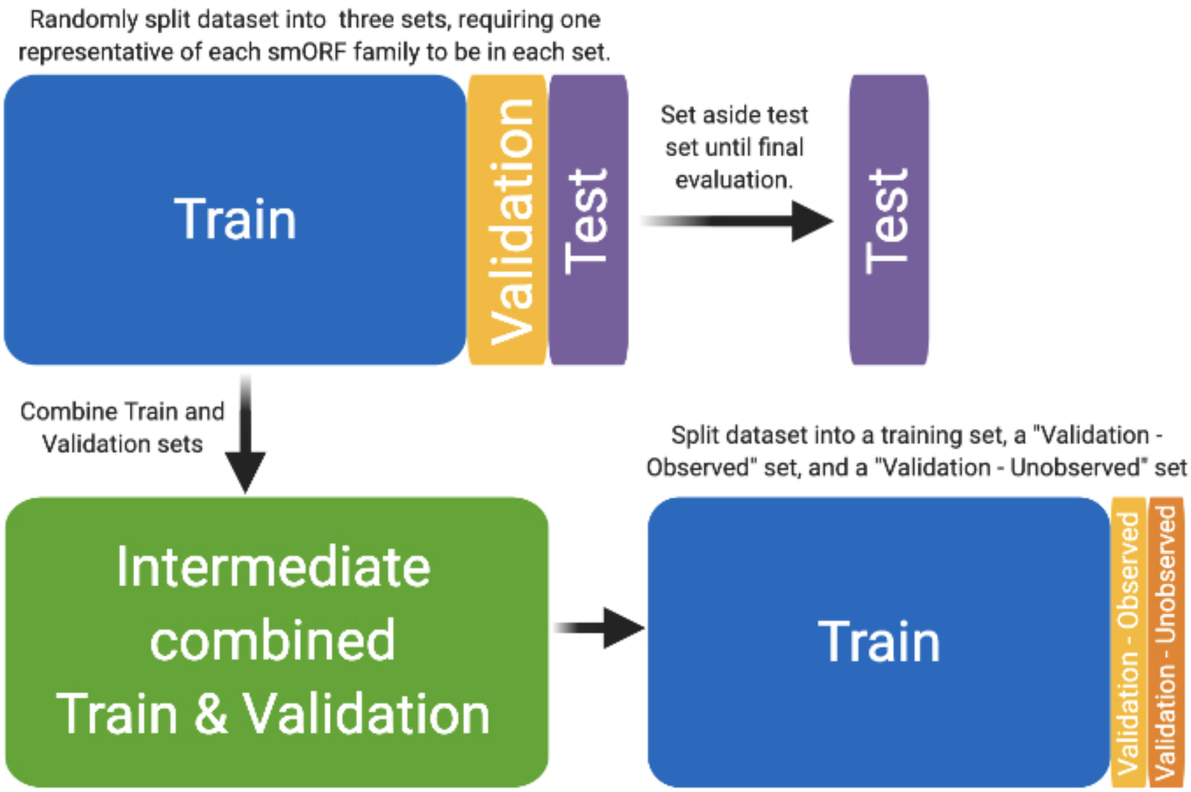


**Figure S1. Schematic of deep learning validation and evaluation strategy.** This schematic illustrates how the full dataset was split into training, validation, and test sets. First, the full dataset was split into training, validation, and test. Each division contains at least one representative of each smORF family. The rest of the examples were randomly assigned to each division, with approximately 80% being used as training, 10% for validation, and 10% for test. The test set was set aside and was not used at all for any of model architecture selection and hyperparameter tuning. The training and validation sets were recombined, and reallocated in such a way that the validation set was approximately 50% unobserved smORF families (not found in the training set) and 50% observed smORF families. Model architecture selection and hyperparameter tuning was done on this dataset, and performance in both the “Validation - Observed” and “Validation - Unobserved” datasets were considered when selecting the final DSN1 and DSN2 models, respectively.

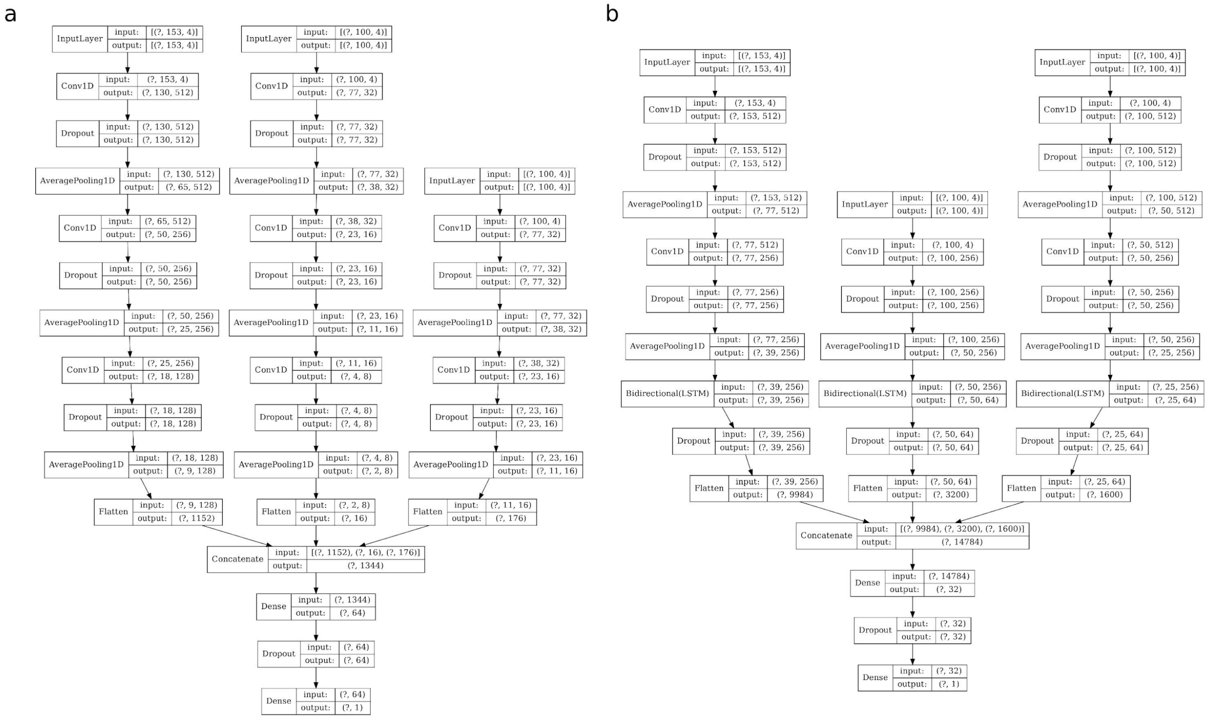


**Figure S2. Schematic of deep learning architectures of DSN1 and DSN2.** Schematics representing the model architectures of the DSN1 (a) and DSN2 (b) deep learning models. These were the final models chosen after an extensive hyperparameter tuning process. Each candidate smORF is broken up into three parts - the smORF sequence itself (leftmost branch in each model), 100 nucleotides upstream of the smORF (the middle branch), and 100 nucleotides downstream of the smORF. The input and output dimensions of each matrix in each layer is given. a) The DSN1 model includes three convolutional layers to process the smORF sequence, with 512, 256, and 128 filters per layer of sizes 24, 16, and 8, respectively. The upstream sequence is processed by three convolutional layers, with 32, 16, and 8 filters per layer of sizes 24, 16, and 8, respectively. The downstream sequence is processed by two convolutional layers, with 32 and 16 filters per layer of sizes 24 and 16, respectively. The dropout rate used to regularize the weights of convolutional layers was 0.5, “valid” padding was used with each layer, and average pooling was used after each convolutional layer. The final dense layer that analyzes the flattened and concatenated output of all three branches includes 64 neurons, and a dropout rate of 0.5. Adam optimization with a learning rate of 1e-4 was used to train the model (Kingma & Ba, 2014). b) The DSN2 model includes two convolutional layers to process the smORF sequence, with 512 and 256 filters per layer of sizes 6 and 4, respectively, and a long short-term memory (LSTM) layer containing 128 neurons. The upstream sequence is processed by one convolutional layer with 256 filters of length 12, and an LSTM layer containing 32 neurons. The downstream sequence is processed by two convolutional layers with 512 and 256 filters per layer of sizes 24 and 16, respectively, and an LSTM layer containing 32 neurons.

The dropout rate used to regularize the weights of convolutional layers was 0.5, the dropout rate for the LSTM layers was 0.3, “same” padding was used with each layer, and average pooling was used after each convolutional layer. The final dense layer that analyzes the flattened and concatenated output of all three branches includes 32 neurons, and a dropout rate of 0.5. Adam optimization with a learning rate of 1e-3 was used to train the model.

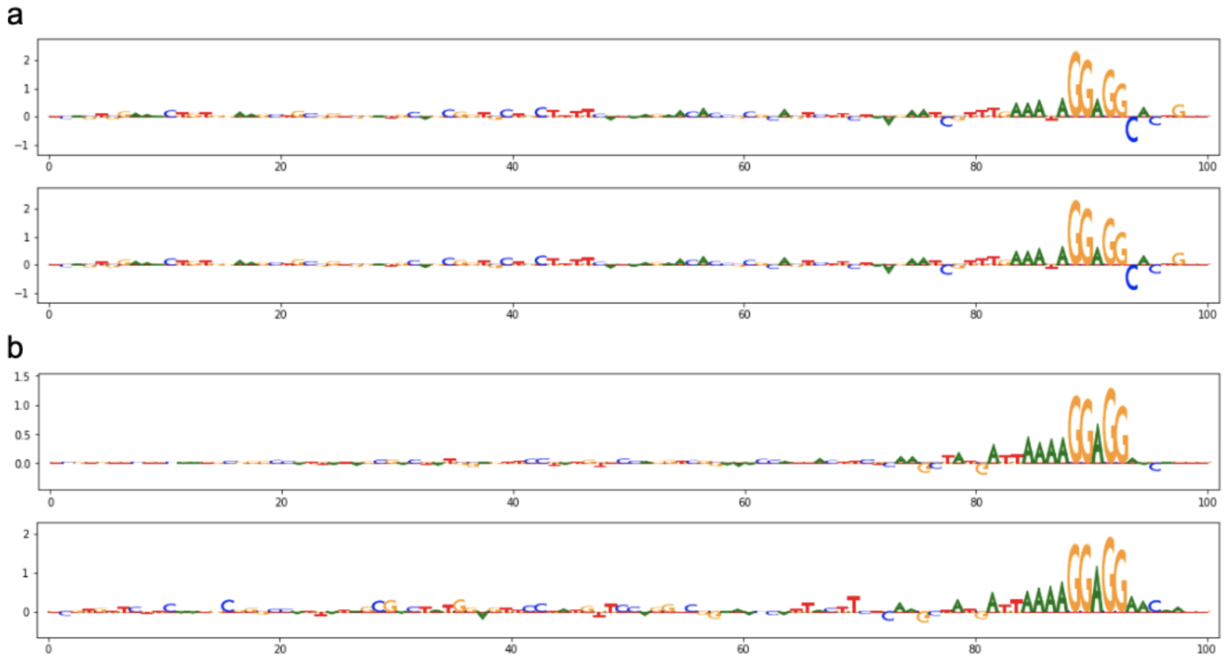


**Figure S3. Examples of Shine-Dalgarno motif identification DSN1 and DSN2 models.** This shows the feature importance scores of upstream sequences as calculated using the DeepLIFT algorithm for two specific examples. This was implemented in the SHAP python package. The canonical AGGAGG shine-dalgarno motif is recognizable, with high importance being assigned to the two guanine repeats. a) Feature importance scores of the upstream sequence of a single smorfam02316 example as assigned by DSN1 (top) and DSN2 (bottom). b) Feature importance scores of the upstream sequence of a single smorfam03028 example as assigned by DSN1 (top) and DSN2 (bottom).

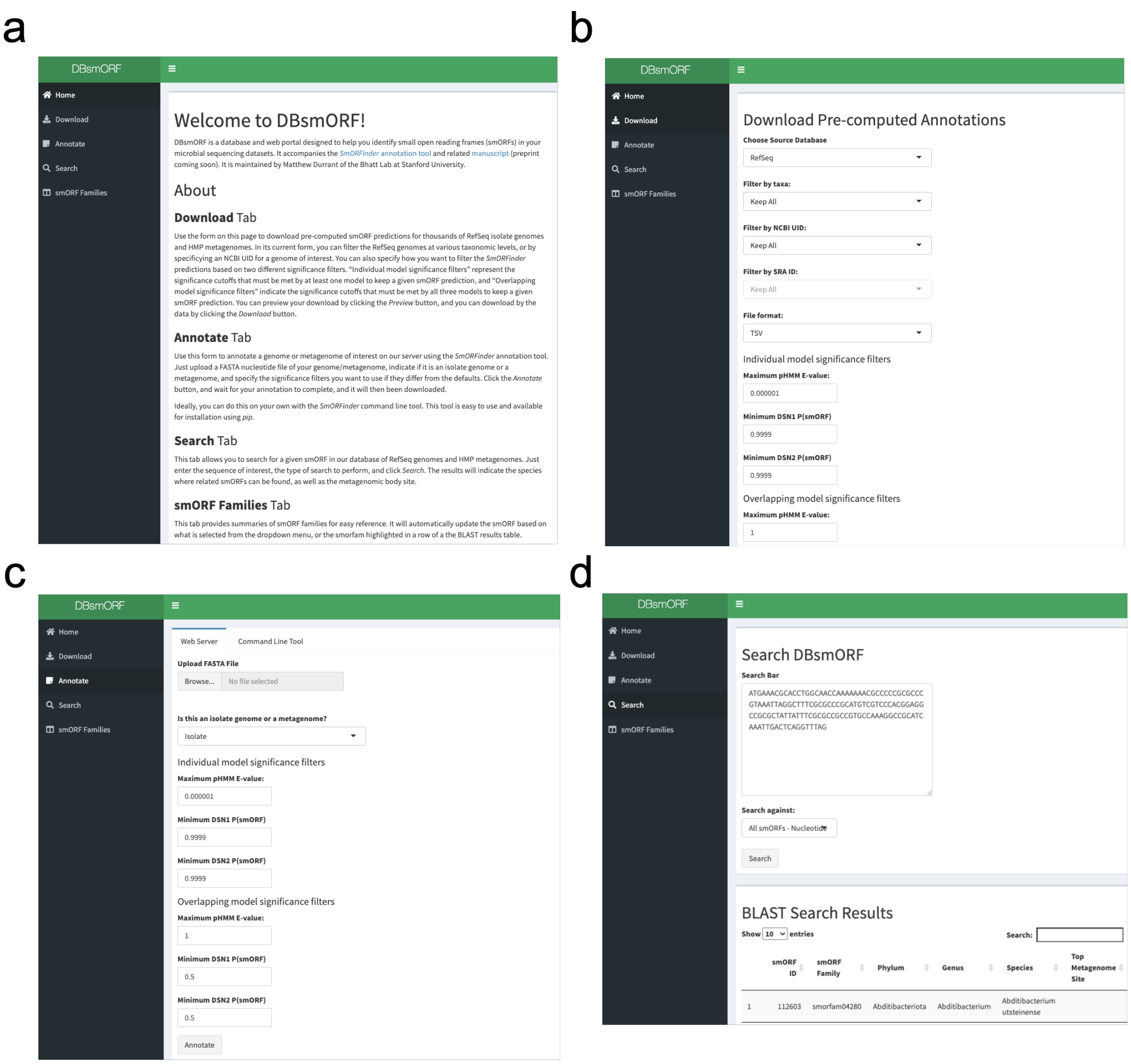


**Figure S4. Web portal for analyzing and annotating smORFs.** Screenshots of the *DBsmORF* web portal that has been made available to the public. Showing the main page (a), a page that allows users to download pre-computed smORF annotations for hundreds of thousands of RefSeq bacterial genomes (b), a page for uploading a genome/metagenome of interest to be annotated using the *SmORFinder* tool (c), and a page where users can run BLAST searches of their smORFs of interest against the database (d).
